# PsiConnect: A Multimodal Neuroimaging Study of Psilocybin-Induced Changes in Brain and Behaviour

**DOI:** 10.1101/2025.04.11.643415

**Authors:** Leonardo Novelli, Devon Stoliker, Tamrin Barta, Matthew D. Greaves, Sidhant Chopra, James Jackson, Jessica Kwee, Martin L. Williams, Adeel Razi

**Affiliations:** School of Psychological Sciences and Monash Biomedical Imaging, Monash University, Australia; Orygen, The University of Melbourne, Parkville, Australia; Centre for Youth Mental Health, The University of Melbourne, Australia; School of Health Sciences, Swinburne University of Technology, Hawthorn, Australia; Wellcome Centre for Human Neuroimaging, University College London, United Kingdom; CIFAR Azrieli Global Scholars Program, Toronto, Canada

## Abstract

PsiConnect is a large-scale neuroimaging study designed to investigate the neural and subjective effects of psilocybin using multimodal neuroimaging. It combines functional, structural, and diffusion-weighted MRI with EEG to examine brain activity in 62 participants before and after a 19 mg dose of psilocybin. The design includes resting-state scans and three naturalistic conditions: guided meditation, music listening, and movie watching. Half of the cohort underwent an 8-week meditation training program, enabling the exploration of interactions among meditation, psilocybin, and brain function. The fMRI data was obtained through multi-echo fMRI, which enhances the signal-to-noise ratio and reduces susceptibility artifacts, thereby improving the reliability of the analyses. A comprehensive battery of behavioural and self-report measures captured both acute and longitudinal cognitive and subjective effects, with follow-ups extending to one year post-administration. The large sample size, multimodal neuroimaging, diversity of contexts, and longitudinal behavioural follow-ups enable the study of psilocybin-induced changes in brain and behaviour with an unprecedented level of detail and reliability. Furthermore, the data is curated according to open science principles to ensure accessibility and interoperability with established neuroimaging processing pipelines. These factors make PsiConnect a valuable and highly reusable resource for researchers in cognitive and computational neuroscience.

## Background & Summary

PsiConnect (a portmanteau of Psilocybin, Connectivity, and Context) is a large-scale neuroimaging study examining psilocybin’s effects on the human brain. Conducted at Monash University, it provides the world’s largest single-site neuroimaging dataset on serotonergic psychedelics, with 62 participants scanned before and after receiving a 19 mg oral dose of psilocybin. The study is multimodal, involving functional, anatomical, and diffusion MRI, as well as EEG recordings, to deepen our understanding of psilocybin’s effects at multiple neurophysiological scales. Both MRI and EEG scans included resting-state and three naturalistic stimuli: guided meditation, music listening, and movie watching. These enable the study of how psilocybin influences both spontaneous and stimulus-driven brain activity. Additionally, half of the participants completed an 8-week meditation training program, providing the opportunity to investigate the interactions between meditation, psilocybin, and brain function. The neuroimaging data is complemented by an extensive set of behavioural questionnaires that capture subjective experiences and cognitive changes induced by psilocybin. Participants were assessed longitudinally, with follow-ups extending up to one year post-administration, enabling the study of both acute and sustained changes. These assessments include self-report measures of mental wellbeing, personality traits, and psychedelic experiences, allowing for correlations between subjective reports and neuroimaging findings. The breadth and depth of the behavioural data further enhance the reuse value of the dataset, providing opportunities for interdisciplinary research in psychology and neuroscience. The study design is illustrated in Fig. 1.

**Figure 1.**
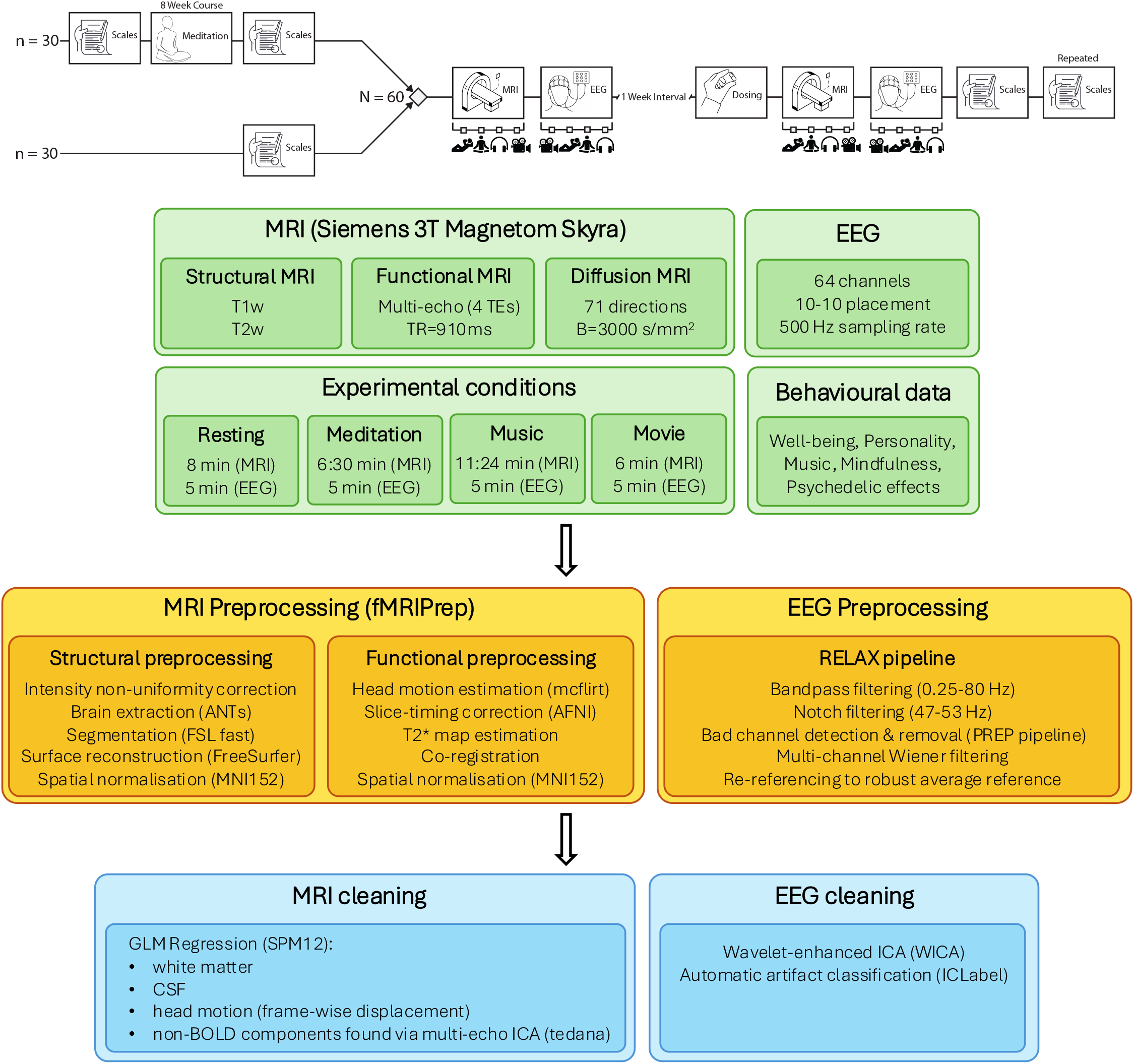
**a)** Simplified representation of study design. Participants were assigned to 8 week mindfulness meditation course or immediately enrolled to multimodal (fMRI and EEG) imaging sessions. An open label imaging session was conducted approximately one week before imaging under administration of 19 mg psilocybin. Music, mindfulness, personality and psychedelic measures were collected before and longitudinally after psilocybin administration. Meditators received additional assessments before and after the mindfulness course. **b)** Data acquisition, preprocessing, and cleaning pipelines.

A key methodological feature of PsiConnect is the use of multi-echo fMRI, which enhances signal-to-noise ratio without sacrificing temporal resolution. Multi-echo acquisition generates multiple images per volume at different echo times spanning tens of milliseconds, covering the entire range of signal decay rates in different voxels across the brain. While this comes at the expense of additional complexity in acquisition and pre-processing and reduced spatial resolution, it offers several advantages: 1) The echoes can be optimally weighted and combined into a single time series to increase the BOLD signal-to-noise ratio, 2) reduced signal dropout due to susceptibility artifacts, and 3) improved test-retest reliability of resting-state functional connectivity^1,2^. Furthermore, multi-echo independent component analysis (ME-ICA) can identify nuisance signal components that do not exhibit typical BOLD signal decay over echo times^3^, which can enhance denoising compared to single-echo acquisitions. Finally, effect size estimates can increase by more than 50% in brain regions prone to signal dropout^4^. The combination of high temporal resolution, robust acquisition methods, and sophisticated preprocessing techniques ensures that the dataset is well-suited for studies on functional connectivity, brain network dynamics, and computational modelling.

PsiConnect is the only fMRI study of psilocybin to combine resting-state scans with three distinct naturalistic stimuli. It is complementary to the recent dataset by Siegel et al.^5^, which emphasizes precision functional mapping with multiple time points and an active placebo condition, but has a sample size of 7 participants. While Siegel et al. provide a longitudinal perspective, PsiConnect offers the largest sample size to date and enables the study of psilocybin’s effects across diverse contexts. By incorporating naturalistic conditions that closely align with practices in clinical research and emerging therapeutic applications, PsiConnect provides a unique opportunity to examine psilocybin’s impact on large-scale brain dynamics in ecologically valid contexts.

The study design and methodology ensure broad applicability across cognitive neuroscience, clinical research, and computational modelling, enabling researchers across various disciplines to leverage PsiConnect to examine fundamental questions about subjective experience, brain connectivity, and altered states of consciousness. Finally, all data is open-access via OpenNeuro. For these reasons, PsiConnect has high reuse potential for investigating psychedelic effects on the brain and behaviour.

Here, we introduce the PsiConnect dataset structure and neuroimaging acquisition protocols. We detail the organisation of neuroimaging and behavioural data according to open science best practices to facilitate interoperability with existing software and processing pipelines. Researchers can use this information to process the raw data according to the specific needs of their methods. For those who wish to use the data seamlessly without reprocessing it, we release the minimally preprocessed data and analysis-ready cleaned data, explaining the processing steps in detail and providing quality control assessments to evaluate the efficacy of the preprocessing pipeline. Our extensive quality control assessments confirms the high quality of the data and its value for future research.

## Methods

### Ethics and clinical trial registration

The study protocol was approved by the Monash University Human Research Ethics Committee. The trial was registered with the Australian New Zealand Clinical Trials Registry under the registration number AC-TRN12621001375842.

### Study design

The study was open-label and included two imaging sessions: a baseline (no-psilocybin) session and a session following the administration of a 19 mg dose of psilocybin (see Fig. 1). Both sessions involved MRI and EEG scans, with four conditions repeated in each: resting state, guided meditation, music listening, and movie watching. In the resting-state condition (8:00 minutes), participants were instructed to relax and keep their eyes closed while remaining still. During the guided meditation (6:30 minutes), participants received brief instructions via MRI-safe audio, followed by silent periods of meditation. For the music listening condition (11:24 minutes), a curated playlist was designed to evoke emotional depth and resonance. In the naturalistic movie condition (6:00 minutes), participants watched a video of clouds moving at a natural (imperceptible) pace, without audio. These conditions were studied in both the baseline (no-psilocybin) and psilocybin sessions and were repeated in both fMRI and EEG, allowing for comprehensive cross-modal analyses of psilocybin’s influence on brain activity and connectivity. The eligible participants were stratified based on age and sex and then randomly assigned to one of the groups. Half of the participants were assigned to an 8-week mindfulness-based cognitive therapy (MBCT) program *Finding Peace in a Frantic World* run by a trained and registered instructor, which involved weekly online group classes and daily independent practice; the other half were assigned to a control group with no intervention. Several behavioural measures were collected before and during the baseline (no-psilocybin) and psilocybin scans (see *Behavioural Data Acquisition*). The follow-up conducted the day after psilocybin administration included semi-structured, open-ended questions and experience ratings. Further follow-up measures were administered one week, and one, three, six and twelve months after psilocybin administration.

### Participants

Sixty-five healthy, psychedelic-naive adults aged 18-55 were recruited for the study. Participants were required to have no formal meditation practice and limited previous exposure to meditation. They were first screened via short online survey and then detailed screening was performed by a suitably trained staff member for excluding any psychopathology using the long form SCID-V^6^. Exclusion criteria included a history of psychiatric disorders or suicidality, a 5-year history of substance and/or alcohol use disorder, first-degree relatives with a diagnosed psychotic disorder, a history of major neurological disorders including stroke and epilepsy, use of hallucinogens within the last six months, a formal meditation practice in the last 6 months or extensive previous exposure to mindfulness mediation, and contraindicated medications. Participants were also screened for MR contraindications and provided informed consent. On the day of psilocybin administration, they were assisted by a study doctor, researchers, lab staff with relevant training, and volunteers from the community drug harm reduction support organisation *Dancewize*. Adverse effects following psilocybin administration were rare and generally mild. Three participants reported transient headaches during the night (n=3). Additionally, three participants required follow-up support from a clinical psychologist familiar with psychedelic integration. These follow-up calls were conducted over the phone, and no further assistance was required. Two participants didn’t complete the psilocybin imaging session (neither MRI nor EEG, due to a complaint of back-pain which subsided once psilocybin effects also subsided, and an incidental finding on their baseline scan in the other), and one only completed part of the resting-state MRI scan due to adverse responses (a combination of dysphoria, panic, and distressing closed-eye visuals in the MRI). Two others didn’t complete the second EEG session due to delayed onset of feeling overwhelmed, and one was excluded because they disclosed having extensive meditation experience at the time of baseline imaging, which was an exclusion criteria. However, on no ocassion it was deemed necesssary by the study doctor to adminster an anti-depresssant. This means that fMRI raw data is available for sixty-four participants in the baseline session, and sixty-one participants in the psilocybin session. EEG raw data is available for sixty-four participants in the baseline session, and fifty-nine participants in the psilocybin session.

### Data acquisition

#### Structural MRI acquisition

Structural MRI data was acquired using a Siemens 3 Tesla Magnetom Skyra scanner at Monash Biomedical Imaging, Monash University, Australia. T1-weighted (T1w) anatomical images were obtained for each participant during two sessions: at baseline (no-psilocybin) and on the psilocybin administration day. The images were acquired using a 3D magnetisation-prepared rapid gradient-echo (MP-RAGE) sequence with a 32-channel head coil. The acquisition parameters were as follows: repetition time (TR) of 2300 ms, echo time (TE) of 2.07 ms, 192 slices per slab, 1 mm slice thickness, and 1 mm isotropic voxel size. The parallel acquisition technique mode used was GRAPPA, with an acceleration factor (PE) of 2 on the baseline day and 3 on the psilocybin administration day. A T2-weighted anatomical image was obtained only during the baseline session. The acquisition parameters were: TR of 3200 ms, TE of 452 ms, 176 slices per slab, 1 mm slice thickness, and 1 mm isotropic voxel size, with an acceleration factor of 2.

#### Functional MRI acquisition

Blood-oxygenation-level-dependent (BOLD) fMRI data was collected using a multi-echo, multi-band, echo-planar imaging, T2*-weighted sequence. The acquisition parameters were: TR of 910 ms, multi-echo TE of 12.60 ms, 29.23 ms, 45.86 ms, 62.49 ms, multi-band acceleration factor of 4, field of view of 206 mm, RL phase encoding direction, and 3.2 mm isotropic voxels. The scan durations were: resting state with eyes closed (8 minutes, 505 volumes), audio-guided meditation with eyes closed (6:30 minutes, 405 volumes), music listening with eyes closed (11:24 minutes, 728 volumes), movie watching (6:00 minutes, 372 volumes). The structural and functional MRI images acquired from the Siemens scanner were converted into the Neuroimaging Informatics Technology Initiative (NIfTI) format.

#### Diffusion Weighted Imaging Acquisition

Diffusion weighted imaging data were acquired using a single-shell acquisition scheme with an echo-planar imaging sequence. The acquisition parameters were as follows: TR of 5900 ms, TE of 171.0 ms, 56 slices per slab, 2.5 mm slice thickness, and 2.5 mm isotropic voxel size. The field of view was 212 mm in the read direction and 100% in the phase direction. The Parallel Acquisition Technique mode used was GRAPPA, with an acceleration factor (PE) of 2 and 40 reference lines in the phase-encoding direction. Diffusion weighting was applied along 71 directions using a monopolar diffusion scheme. Two b-values were used: 0 s/mm^2^ and 3000 s/mm^2^. The EPI factor was set to 86, and the bandwidth was 1038 Hz/Px. Fat suppression was implemented using a fat saturation technique.

#### EEG acquisition

EEG data was recorded using a 64-channel BrainAmp MR Plus amplifier (Brain Products GmbH, Germany) with BrainVision Recorder software (version 1.22.0001, Brain Products GmbH, Germany) and Ag/AgCl electrodes embedded in an actiCAP slim cap according to the standardized 10-10 system^7^. The FCz electrode served as the reference and the FPz electrode served as the ground for online recording. The impedances were maintained below 10*k*Ω using an abrasive paste (Nuprep, Weaver and Company, USA) followed by applying a conductive gel (Easycap GmbH, Germany). EEG signals were sampled at 500 Hz with a bandpass filter of 0.01 *−* 1000*Hz*. Participants were comfortably seated in an acoustically absorbent room to minimise environmental noise and their chin strap was securely fastened to ensure head stability. They were instructed to remain as still as possible and avoid jaw clenching during the recording to minimise artifacts.

The EEG recording session included the same four conditions as the fMRI described above. The naturalistic movie (5 minutes) was presented first, followed by the three eyes-closed conditions: resting state (5 minutes), audio-guided meditation (5 minutes), and music listening (7 minutes). The raw data was acquired as a single continuous recording over all conditions and then split into four separate files using the event markers for the start and end of each condition. Due to technical issues, event markers were not recorded for two participants in the baseline session and one participant in the psilocybin session, so these recordings were excluded from pre-processing and analysis. In summary, pre-processed EEG is available for 63 participants in the baseline session and 58 in the psilocybin session.

#### Behavioural data acquisition

A synopsis of the behavioural measures and delivery times is provided in Tables 2 and 3. Survey presentation was randomised between participants, whereby all non-mood related measures were presented in a random order first, excluding the DASS-21 and IDAS-II. Both these scales always appeared at the end, and were randomised within each other. Participants in the meditation arm completed the following measures prior to meditation training: Depression Anxiety and Stress Scale (DASS-21), selected scales from an abridged version of the Expanded Version of the Inventory of Depression and Anxiety Symptoms (IDAS-II), The Warwick-Edinburgh Mental Wellbeing Scale (WEMWBS), and State Adult Attachment Measure (SAAM). Each week of the meditation course, participants were instructed to complete the Mindfulness Adherence Questionnaire (MAQ). They also completed additional questions about their meditation practice, as described in the metadata (JSON) files. Prior to their baseline scanning sessions, all participants completed the following measures at home: Political Perspective Questionnaire (PPQ-5), Nature Relatedness Scale (NR-6), The Attachment Style Questionnaire (ASQ), Levenson Multidimensional Locus of Control Scales (LOC), NEO Five-Factor Inventory-3 (NEO-FFI-3), Aesthetic Experiences Scale in Music (AES-M), Absorption in Music Scale (AIMS), Barcelona Music Reward Questionnaire (BMRQ), Short Test of Musical Preferences (STOMP-R), and Life Attitude Profile-Revised (LAP-R).

On the baseline scan day, participants completed several measures after the MRI. They first completed a measure evaluating perceptions of patterns, faces, and imagery observed during the video of clouds (referred to as “clouds” or “clouds measure”). This was followed by a measure assessing ratings of liking, resonance, and openness towards the music, referred to here as the Music Experience Scale (MES). The order of these measures was kept consistent. During EEG setup, participants listened to five music samples, rating each using the MES before advancing to the next sample. The order of the samples was randomised. The first EEG recording was the video of clouds. Following the video, participants completed the clouds measure. After this, EEG recording continued with rest, meditation, and music conditions. After EEG recording was completed, they completed the MES. Shortly after, participants completed a series of psychological measures, including the Five Facet Mindfulness Questionnaire (FFMQ-SF), DASS-21, IDAS-II, WEMWBS, and the SAAM. Additionally, a 42-item version of the 5-Dimension Altered States of Consciousness (5D-ASC) scale was administered, allowing for the derivation of the 11-Dimension ASC (11D-ASC) solution, as described in^8^. Participants in the meditation condition also completed the Meditation Depth Questionnaire (MEDEQ)^9^.

On the administration day, participants again first completed the MES and clouds measure after the MRI, followed by the Ego Dissolution Inventory (EDI). The order of these measures was kept consistent. During EEG setup, participants listened to five music samples, rating each using the MES before advancing to the next sample. The order of the samples was randomised. The first EEG recording was the video of clouds. Following the video, participants completed the clouds measure. After this, EEG recording continued with rest, meditation, and music conditions. After the completion of EEG recording, the MES was again administered. At the end of the session, participants completed the 94-item 5D-ASC (with 94 items used to construct the 5D-ASC and 42 items used to construct the 11D-ASC), the Mystical Experience Questionnaire - Revised (MEQ30), the AES-M, and a second administration of the EDI. The order of the measures was randomised. The metadata (JSON) files for all sessions provide the questions asked and more details.

The one-day follow-up included semi-structured, open-ended questions, experience ratings, and meditation evaluations for the meditation arm. Ratings of personal meaningfulness and spiritual significance^10^, subjective intensity, and anxiety at times throughout the day were also included. Further follow-up measures were administered one week, and one, three, six and twelve months after psilocybin administration. At each follow-up time point, participants completed the Self-Compassion Scale (SCS), FFMQ-SF, and meditation participants additionally completed the MAQ. The one-week follow-up also included the Life Changes Inventory – Revised, AES-M, Emotional Breakthrough Inventory (EBI), LOC, and SAAM. The one-month follow-up included: Persisting Effects Questionnaire Revised (PEQ60), Experiences Questionnaire (EQ), LAP-R, PPQ-5, NR-6, ASQ, LOC, DASS-21, IDAS-II, and meditation participants completed the Meditation Depth Questionnaire (MEDEQ) instead of the MAQ. Finally, the 12-month follow-up also included the LAP-R instead of the LOC and both the PPQ-5 and NR-6.

### Data preprocessing and cleaning

#### BIDS

The Digital Imaging and Communications in Medicine (DICOM) images acquired from the Siemens (Skyra) scanner were converted into the NIfTI format and organized according to the Brain Imaging Data Structure (BIDS) 1.7.0 using BIDScoin^11^. Following the BIDS guidelines, each data file is accompanied by a corresponding metadata file in JSON format. These contain extensive lists of neuroimaging acquisition parameters, and the necessary information to interpret the behavioural measures. To de-identify the anatomical images, the facial features were removed using PyDeface (https://github.com/poldracklab/pydeface). The EEG data was also organised according to BIDS 1.7.0.

#### MRI anatomical data preprocessing

Anatomical MRI preprocessing was performed using *fMRIPrep* 22.0.2^12,13^(RRID:SCR_016216), which is based on *Nipype* 1.8.5^14,15^ (RRID:SCR_002502). The baseline and administration T1-weighted (T1w) images were corrected for intensity non-uniformity (INU) with `N4 BiasFieldCorrection`^16^, distributed with ANTs 2.3.3^17^ (RRID:SCR_004757). The T1w-reference was then skull-stripped with a *Nipype* implementation of the `antsBrainExtraction.sh` workflow (from ANTs), using OASIS30ANTs as the target template. Brain tissue segmentation of cerebrospinal fluid (CSF), white matter (WM) and gray matter (GM) was performed on the brain-extracted T1w using `fast`^18^ (FSL 6.0.5.1:57b01774, RRID:SCR_002823). A T1w-reference map was computed after registration of 2 T1w images (after INU-correction) using `mri_robust_template`^19^ (FreeSurfer 7.2.0). Brain surfaces were reconstructed using `recon-all`^20^ (FreeSurfer 7.2.0, RRID:SCR_001847), and the brain mask estimated previously was refined with a custom variation of the method to reconcile ANTs-derived and FreeSurfer-derived segmentations of the cortical gray-matter of Mindboggle^21^ (RRID:SCR_002438). Volume-based spatial normalization to one standard space (MNI152NLin2009cAsym) was performed through nonlinear registration with `antsRegistration` (ANTs 2.3.3), using brain-extracted versions of both T1w reference and the T1w template. The following template was selected for spatial normalization: *ICBM 152 Nonlinear Asymmetrical template version 2009c*^22^ (RRID:SCR_008796; TemplateFlow ID: MNI152NLin2009cAsym).

#### MRI functional data preprocessing

Functional MRI preprocessing was also performed using *fMRIPrep* 22.0.2^12,13^. For each of the 8 BOLD runs per participant (across all tasks and sessions), the following preprocessing was performed. First, a reference volume and its skull-stripped version were generated from the shortest echo of the BOLD run using a custom methodology of *fMRIPrep*. Head-motion parameters with respect to the BOLD reference (transformation matrices, and six corresponding rotation and translation parameters) were estimated before any spatiotemporal filtering using `mcflirt`^23^ (FSL 6.0.5.1:57b01774). BOLD runs were slice-time corrected to 0.401 s (0.5 of slice acquisition range 0s-0.802 s) using `3dTshift` from AFNI^24^ (RRID:SCR_005927). The BOLD time-series (including slice-timing correction) were resampled onto their original, native space by applying the transforms to correct for head-motion. These resampled BOLD time-series will be referred to as *preprocessed BOLD in original space*, or just *preprocessed BOLD*. A T2^⋆^ map was estimated from the preprocessed EPI echoes, by voxel-wise fitting the maximal number of echoes with reliable signal in that voxel to a monoexponential signal decay model with nonlinear regression. The T2^⋆^/S0 estimates from a log-linear regression fit were used for initial values. The calculated T2^⋆^ map was then used to optimally combine preprocessed BOLD across echoes following the method described in^25^. The optimally combined time series was carried forward as the *preprocessed BOLD*. The BOLD reference was then co-registered to the T1w reference using `bbregister` (FreeSurfer) which implements boundary-based registration^26^. Co-registration was configured with six degrees of freedom. Several confounding time series were calculated based on the *preprocessed BOLD*: framewise displacement (FD), DVARS and three region-wise global signals. FD was computed using two formulations following Power (absolute sum of relative motions,^27^) and Jenkinson recommendations (relative root mean square displacement between affines,^23^). FD and DVARS are calculated for each functional run, both using their implementations in *Nipype*, following the definitions by^27^. The three global signals are extracted within the CSF, the WM, and the whole-brain masks. The BOLD time series were resampled in standard space, generating a *preprocessed BOLD run in MNI152NLin2009cAsym space*. All resamplings were performed with *a single interpolation step* by composing all the pertinent transformations (i.e. head-motion transform matrices and co-registrations to anatomical and output spaces). Gridded (volumetric) resamplings were performed using `antsApplyTransforms` (ANTs), configured with Lanczos interpolation to minimize the smoothing effects of other kernels^28^. Non-gridded (surface) resamplings were performed using `mri_vol2surf` (FreeSurfer). Many internal operations of *fMRIPrep* use *Nilearn* 0.9.1^29^ (RRID:SCR_001362), mostly within the functional processing workflow. For more details of the pipeline, see the section corresponding to workflows in *fMRIPrep*’s documentation.

#### Additional available regressors

Additionally, a set of physiological regressors were extracted to allow for component-based noise correction (*CompCor*^30^). Principal components are estimated after high-pass filtering the *preprocessed BOLD* time series (using a discrete cosine filter with 128 s cut-off) for the two *CompCor* variants: temporal (tCompCor) and anatomical (aCompCor). tCompCor components are then calculated from the top 2% variable voxels within the brain mask. For aCompCor, three probabilistic masks (CSF, WM and combined CSF+WM) are generated in anatomical space. The implementation differs from that of Behzadi et al. in that instead of eroding the masks by 2 pixels on BOLD space, a mask of pixels that likely contain a volume fraction of GM is subtracted from the aCompCor masks. This mask is obtained by dilating a GM mask extracted from the FreeSurfer’s *aseg* segmentation, and it ensures components are not extracted from voxels containing a minimal fraction of GM. Finally, these masks are resampled into BOLD space and binarized by thresholding at 0.99 (as in the original implementation). Components are also calculated separately within the WM and CSF masks. For each CompCor decomposition, the *k* components with the largest singular values are retained, such that the retained components’ time series are sufficient to explain 50% of variance across the nuisance mask (CSF, WM, combined, or temporal). The remaining components are dropped from consideration.

The head-motion estimates calculated in the correction step were also placed within the corresponding confounds file. The confound time series derived from head motion estimates and global signals were expanded with the inclusion of temporal derivatives and quadratic terms for each^31^. Frames that exceeded a threshold of 0.5 mm FD or 1.5 standardized DVARS were annotated as motion outliers. Additional nuisance time series are calculated by means of principal components analysis of the signal found within a thin band (*crown*) of voxels around the edge of the brain, as proposed by^32^.

#### fMRI data cleaning

The preprocessed and optimally combined data produced by *fMRIPrep* was cleaned via a single regression in *SPM12*^33^ using the general linear model. The regressors were: the white matter and cerebrospinal fluid signals computed by *fMRIPrep*, the frame-wise displacement, and the non-BOLD components identified via multi-echo ICA performed using *tedana* (version 0.0.12)^34^. In short, multi-echo ICA identifies non-BOLD components that are independent of the echo time^3^. Finally, to facilitate surface-based analyses, the cleaned volumetric data was projected to the FreeSurfer left-right-symmetric cortical surface template with 32000 vertices for each hemisphere (fsLR32k).

#### EEG data preprocessing and cleaning

The raw EEG signals were preprocessed using the default parameters of the automated RELAX pipeline^35^ implemented in MATLAB using functions from EEGLAB^36^ and FieldTrip^37^ toolboxes. First, the data was bandpass filtered between 0.25 and 80 Hz using a fourth-order Butterworth filter with zero phase, with a notch filter applied between 47 and 53 Hz to reduce line noise. Subsequently, bad channels were removed by applying a multistep process incorporating the “findNoisyChannels” function of the PREP pipeline^38^. This was followed by the initial reduction of the artefacts related to eye movements, muscle activity and drift using a multi-channel Wiener filter^39^, after which the data were re-referenced to the robust average reference^38^. Residual artefacts were then removed by Wavelet-enhanced Independent Component Analysis (WICA) with the artefactual components identified by the automated ICLabel classifier^40^. Electrodes rejected during the cleaning process were then restored to the data through spherical interpolation. Finally, all pre-processed data were visually inspected for data quality.

#### Behavioural data processing

Questionnaires were administered online using Qualtrics. The raw data from all questionnaires were collected in their original Likert scale format, as defined by the authors of each measure. Raw data spreadsheets are provided for each time point. Subscale scores were computed according to the authors’ guidelines, using the referenced papers listed in the metadata (JSON) file for each time point. If fewer than 50% of the required items for a scale were missing, the missing values were imputed using the median of the available items. Both the imputed and not imputed values are released as part of the dataset.

### Data Records

The data can be accessed from the OpenNeuro public repository (access number: ds006110), organized according to the Brain Imaging Data Structure (BIDS) specification, version 1.7.0^41^. Specifically, the defaced raw data and the MRI and EEG derivatives are organised in the subfolders indicated in Table 1. The MRI derivatives are listed in the same order as they were computed via the preprocessing and cleaning pipeline: cortical surface reconstruction via FreeSurfer, minimal preprocessing via fMRIPrep, multi-echo ICA via tedana, and cleaning via a GLM performed using SPM12. The following sections will provide additional information about naming conventions and available formats for the files contained in the folders listed in Table 1.

**Table 1.**
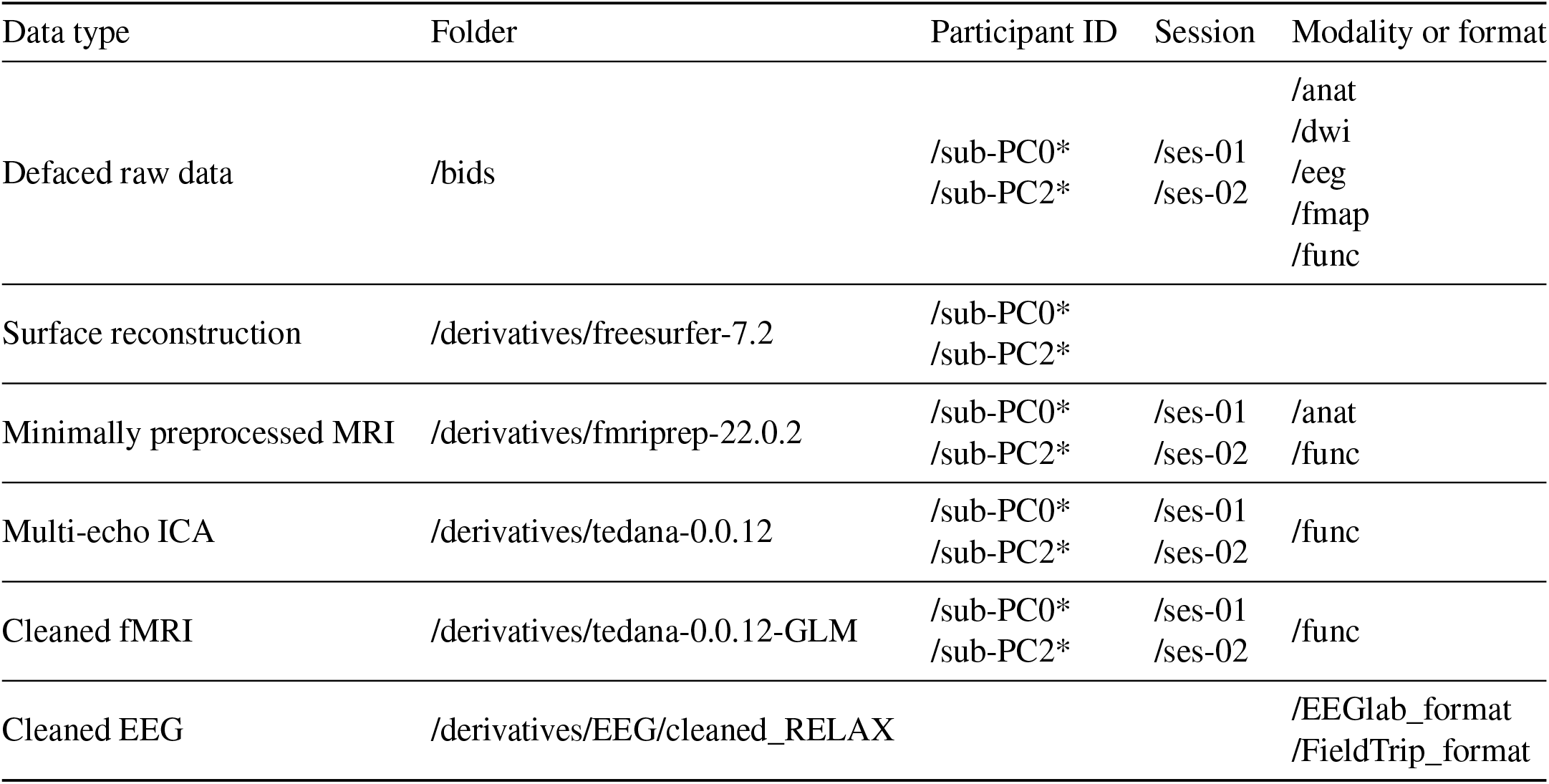
Paths to defaced raw data and derivatives organised according to the Brain Imaging Data Structure (BIDS) 1.7.0. The MRI derivatives are listed in the same order as they were computed via the preprocessing and cleaning pipeline.

#### Raw MRI data organised according to BIDS

De-identified MRI data (with coded subject IDs and defaced structural images) is organised in individual participant directories named “sub-<ID>“. The IDs starting with “PC2” refer to participants who underwent the 8-week meditation training before the neuroimaging sessions. All other IDs start with “PC0”.

#### Preprocessed and cleaned data

Surface reconstruction data is stored in */derivatives/freesurfer-7*.*2*. Minimally preprocessed MRI data is stored in */derivatives/fmriprep-22*.*0*.*2*, which also contains individual participant reports, each named *sub-<ID>*.*html*. The ME-ICA output is stored in */derivatives/tedana-0*.*0*.*12*. For each participant, there is an individual HTML reports, a table listing the accepted and rejected independent components (named *desc-tedana_metrics*.*tsv*), and a single NIFTI file with the BOLD signal obtained by optimally combineing the four echos (file names ending in *optcomb_bold*.*nii*.*gz*). This optimally combined file is cleaned via GLM, and the cleaned data is stored in */derivatives/tedana-0*.*0*.*12-GLM*, with a file name ending in *optcomGLM_bold*.*nii*.*gz*. To facilitate analysis in surface space, this is also projected to the fsLR32k template (name ending in *optcomGLM_fsLR_32k*.*mat*). Cleaned EEG data is stored in */derivatives/EEG/cleaned_RELAX* and is available in both EEGlab and FieldTrip formats.

#### Audio and video stimuli

All audio and video stimuli were prepared and presented using open-source software. To comply with the media formatting requirements of the presentation software PsychoPy3 (version 2021.2.3), used during MRI and EEG scanning, all audio recordings were converted to the WAV format (sampling rate: 44100 Hz; bit depth: 16-bit; stereo), and the silent (clouds) video file was converted to the M4V format with an RGB24 codec (resolution: 1920×1080 pixels; frame rate: 30.00 fps) using functions in VLC media player (version 3.0.12). Audio stimuli were additionally normalized to a peak amplitude of -1.0 dB using functions in Audacity (version 3.1.3). A complete list of stimuli used in MRI and EEG sessions is provided below, including both the calibration audio for volume adjustments and the instructional audio presented prior to the (clouds) video. Due to copyright restrictions, certain audio files are omitted from the repository but are documented here with barcode identifiers available.

### MRI

- Task: Audio calibration Track: India Song (Piano) Author: D’Alessio, C. Duration: 166.58 s Notes: Subject to copyright (and thus omitted from repository). Barcode: 3 149025 048325.
- Task: Meditation Filename (and location): meditation.wav Author: Stoliker, D. Duration: 390.10 s
- Task: Music (part 1) Track: Sunset from the Ethereal Shore (Lauge Rework) Author: Spacecraft, Lauge Duration: 351.56 s Notes: Subject to copyright (and thus omitted from repository).
- Task: Music (part 2) Track: Sollys (Applefish Rework) Author: Lauge, Applefish Duration: 332.38 s Notes: Subject to copyright (and thus omitted from repository).
- Task: Instructions (open eyes and note patterns) Filename (and location): instructions.wav Author: Greaves, M. D. Duration: 18.54 s
- Task: Video Filename (and location): clouds.m4v Author: Stoliker, D. Duration: 360.08 s

### EEG

- Task: Video Filename (and location): clouds_short.m4v Track: N/A Author: Stoliker, D. Duration: 300.00 s
- Task: Meditation Filename (and location): meditation_short.wav Track: N/A Author: Stoliker, D. Duration: 300.00 s
- Task: Music (part 1) Filename (and location): N/A Track: Sunset from the Ethereal Shore (Lauge Rework) Author: Spacecraft, Lauge Abbreviated track duration. Notes: Subject to copyright (and thus omitted from repository).
- Task: Music (part 2) Filename (and location): N/A Track: Sollys (Applefish Rework) Author: Lauge, Applefish Abbreviated track duration. Notes: Subject to copyright (and thus omitted from repository).

#### Behavioural data

Following BIDS conventions, behavioural data is stored in the *phenotype* folder, with subfolders for different survey types: those specific to the meditation arm (pre-training, weekly measures during the 8-week course, and post-training measures collected after both the 8-week course and psilocybin administration), as well as baseline, psilocybin administration, and follow-up surveys (Table 2). Each session folder contains one or more TSV files. Each TSV file refers to one questionnaire and contains the scores for all the scales and subscales involved. Each TSV file is accompanied by a metadata (JSON) file that contains the specific questions asked for each item in the questionnaire. In addition, there is a JSON file for each scale, listed in Table 2 and described in Table 3. Each contains the description of the scale, a reference to the relevant literature, and information about how the items were aggregated into a score for each subscale (for example, by summation, average, or rescaling to the 0-100 range). The raw data is provided in the *phenotype/rawdata* folder, which contains one spreadsheet per questionnaire in xlsx format.

**Table 2.** Pre-scored surveys available for each time point in BIDS format, including acccompanying metadata.

| Survey | Folder | Spreadsheets (TSV format) | Scales (JSON format) |
| --- | --- | --- | --- |
| Meditation | /meditation | meditation-before-training<br>meditation-training<br>meditation-after-training | MAQ<br>MEDITATION |
| No psilocybin | /ses-01 | ses-01<br>NEO-FFI-3<br>BMRQ | AES, ASC11, ASQ,<br>BMRQ, CLOUDS, DASS,<br>EQ, FFMQ, IDAS, LAPR,<br>LOC, MES, MEDEQ, NEO-FFI-3,<br>NR,PPQ, SAAM, WEMWBS |
| Psilocibin | /ses-02 | ses-02 | AES, ASC11, ASC5, CLOUDS,<br>EDI, MES, MEQ30 |
| Follow-up | /followup-1day | followup-1day |  |
|  | /followup-1week | followup-1week | EBI, FFMQ, LCIR, LOC, SAAM |
|  | /followup-1month | followup-1month<br>NEO-FFI-3 | ASQ, DASS, EQ, FFMQ,<br>IDAS, LAPR, LOC, MEDEQ,<br>NR, PEQ60, PPQ, NEO-FFI-3 |
|  | /followup-3month | followup-3month | EBI, FFMQ, LCIR, LOC, SAAM |
|  | /followup-6month | followup-6month | FFMQ, LAPR, LOC, MAQ |
|  | /followup-12month | followup-12month |  |

**Table 3.** Pre and post psilocybin behavioural measures.

| Self-Report Measure | Category | Description | Time of Delivery |
| --- | --- | --- | --- |
| Inventory of Depression and Anxiety Symptoms (IDAS-11) | Mental Well-being | Mood, including depression and well-being over a two-week period | Baseline, 1-month follow-up |
| Depression, Anxiety and Stress Scales (DASS-21) | Mental Well-being | Depression, anxiety and stress symptoms over a one-week period | Baseline, 1-month follow-up |
| Warwick-Edinburgh Mental Well-being Scale (WEMWBS) | Mental Well-being | Mental well-being over a two-week period | Baseline |
| NEO Five-Factor Inventory (NEO-FFI-3) | Personality | Openness, conscientiousness, extraversion, agreeableness, neuroticism | Baseline, 1-month follow-up |
| Political Perspective Questionnaire (PPQ-5) | Personality | Libertarian-authoritarian political perspective | Baseline, 1-month follow-up, 12-month follow-up |
| Nature Relatedness Scale (NR-6) | Personality | Subjective connection to nature | Baseline, 1-month follow-up, 12-month follow-up |
| State Adult Attachment Measure (SAAM) | Personality | Temporary variations in adult attachment | Baseline, 1-week follow-up |
| Attachment Style Questionnaire (ASQ) | Personality | Attachment styles | Baseline, 1-month follow-up |
| Levenson Multidimensional Locus of Control Scales (LOC) | Personality | Three locus of control domains: internality, powerful others, and chance. | Baseline, 1-week follow-up, 1/3/6-month follow-up |
| Life Attitudes Profile - Revised (LAP-R) | Personality | Multidimensional measure of attitudes toward life | Baseline, 1/6/12-month follow-up |
| Absorption in Music Scale (AIMS) | Music | Ability and willingness to allow music to draw them into an emotional experience | Baseline |
| Barcelona Music Reward Questionnaire (BMRQ) | Music | Individual differences in Music Reward experiences | Baseline |
| Short Test Of Music Preferences (STOMP-R) | Music | Music preferences, which are related to personality variables, self-views, and cognitive abilities | Baseline |
| Aesthetic Experiences Scale in Music (AES-M) | Music | Aesthetic experience in music | Baseline, Administration, 1-week follow-up |
| The Five Facet Mindfulness Questionnaire (FFMQ-SF) | Mindfulness | Mindfulness with regards to thoughts, experiences, actions in daily life | Baseline, 1-week follow-up, 1/3/6/12-month follow-up |
| Meditation Depth Questionnaire (MEDEQ) | Mindfulness | Depth of meditative experiences | Baseline, 1-month follow-up |
| Self-Compassion Scale (SCS) | Mindfulness | Capacity for self-compassion | Baseline, 1-week follow-up, 1/3/6/12-month follow-up |
| Experiences Questionnaire (EQ) | Mindfulness | Decentering and rumination | Baseline, 1-month follow-up |
| Mindfulness Adherence Questionnaire (MAQ) + tailored questions | Mindfulness | Quantity and quality of formal and informal mindfulness based meditation practice | Meditation weekly reflections, 3/6/12-month follow-up |
| Emotional Breakthrough Inventory (EBI) | Mental Well-being | Emotional disorder symptoms | 1-week follow-up |
| Persisting Effects Questionnaire (PEQ) | Mental Well-being | Negative emotional states | 1-month follow-up |
| Life Changes Inventory - Revised | Personality | Openness, conscientiousness, extraversion, agreeableness, neuroticism | 1-week follow-up |
| Day-after Qualitative Report (custom design for PsiConnect) | Subjective Experience | Sequential and elaborate description of subjective experience under psilocybin | 1-day post |
| Ego Dissolution Inventory (EDI) | Psychedelic effects | Emotional disorder symptoms | Administration, prior to EEG setup, final measures |
| Visual stimuli task ("Clouds measure") (custom) | Visual effects | Perception of patterns, faces and complex imagery during presentation of naturalistic (experimental) stimuli | Baseline, Administration (after MRI scan, after EEG eyes-open recording) |
| Music experience scale (MES) | Music | Subjective liking, resonance and openness to music | Baseline, Administration, after MRI scan (after each music sample during EEG setup, after EEG recording) |
| Five Dimensions of Altered States of Consciousness (5D-ASC) | Psychedelic effects | Assessment of negative emotional states | Baseline, Administration, final measures |
| Mystical Experiences Questionnaire (MEQ) | Psychedelic effects | Openness, conscientiousness, extraversion, agreeableness, neuroticism | Administration, final measures |

### Technical Validation

#### MRI quality control

Quality control was performed using the MRIQC BIDS app^42^, which uses structural and functional MRI images to compute several quality metrics (see Supplementary Figs. 1 and 2 for a comprehensive overview). These include the temporal signal-to-noise ratio and the frame-wise displacement that quantifies head motion^43^. Results and group reports can be found in the *derivatives/mriqc-22*.*0*.*6* folder. Principal component analysis (PCA) of these QC metrics revealed clusters corresponding to the four echos, and helped identify some outliers in the structural and functional images (see Fig. 2), which were further confirmed by visual inspection. In the case of fMRI, we found that head motion was the dominant factor causing outliers on the PCA plot, as confirmed by the square markers in Fig. 2a that indicate scans with large head motion (mean frame-wise displacement greater than 0.5, which is often used as an exclusion threshold^44^). Indeed, head motion, quantified via the mean framewise displacement, was the feature with the largest contribution to the second principal component (see Supplementary Fig. 3). A detailed analysis of the mean frame-wise displacement showed that participants moved their head more during the psilocybin MRI session than in the previous session without psilocybin (Fig. 2d,b).

**Figure 2.**
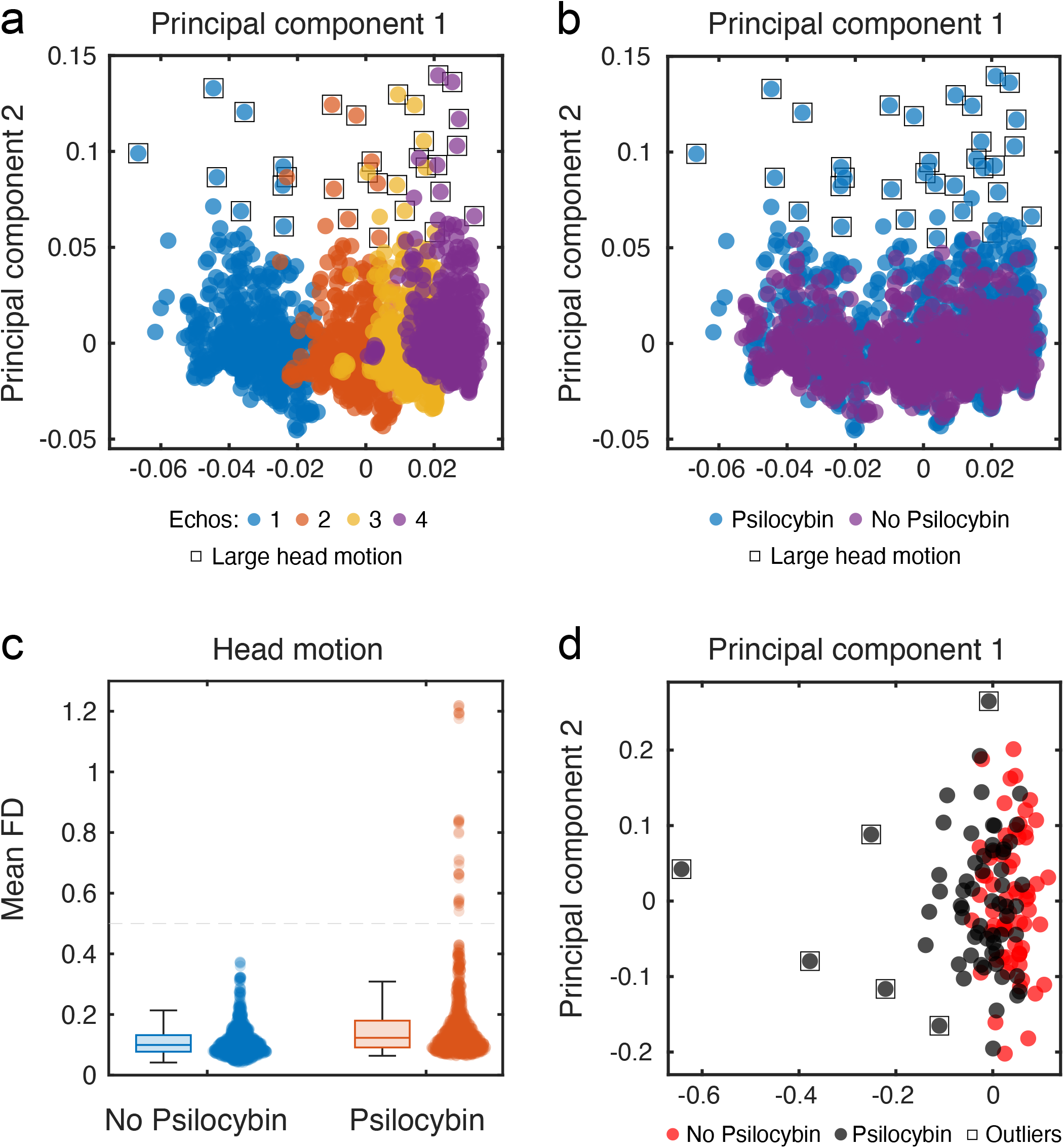
Principal components analysis (PCA) shows that the quality control metrics computed using MRIqc capture important feature of the data and to flag outliers for further inspection. **a)** Functional MRI quality control metrics. Each point represents a single echo, and colour coding the points by echo number showed that PCA was able to cluster the four echos. Points that are further away from the cluster centres are outliers in terms of multiple metrics. However, head motion was the dominant factor in determining these outliers, as shown by the square markers that indicate scans with high head motion (mean frame-wise displacement greater than 0.5). **b)** Colour-coding the same points by session (no psilocybin and psilocybin) shows that the outliers belong to the psilocybin session. **c)** Participants moved their head more during the psilocybin MRI session, in alignment with the findings presented in first two panels. Head motion was quantified using the mean frame-wise displacement (in mm). **d)** Anatomical (T1w) MRI quality control metrics. Square markers indicate outliers that were confirmed by visual inspection of the anatomical scans.

**Figure 3.**
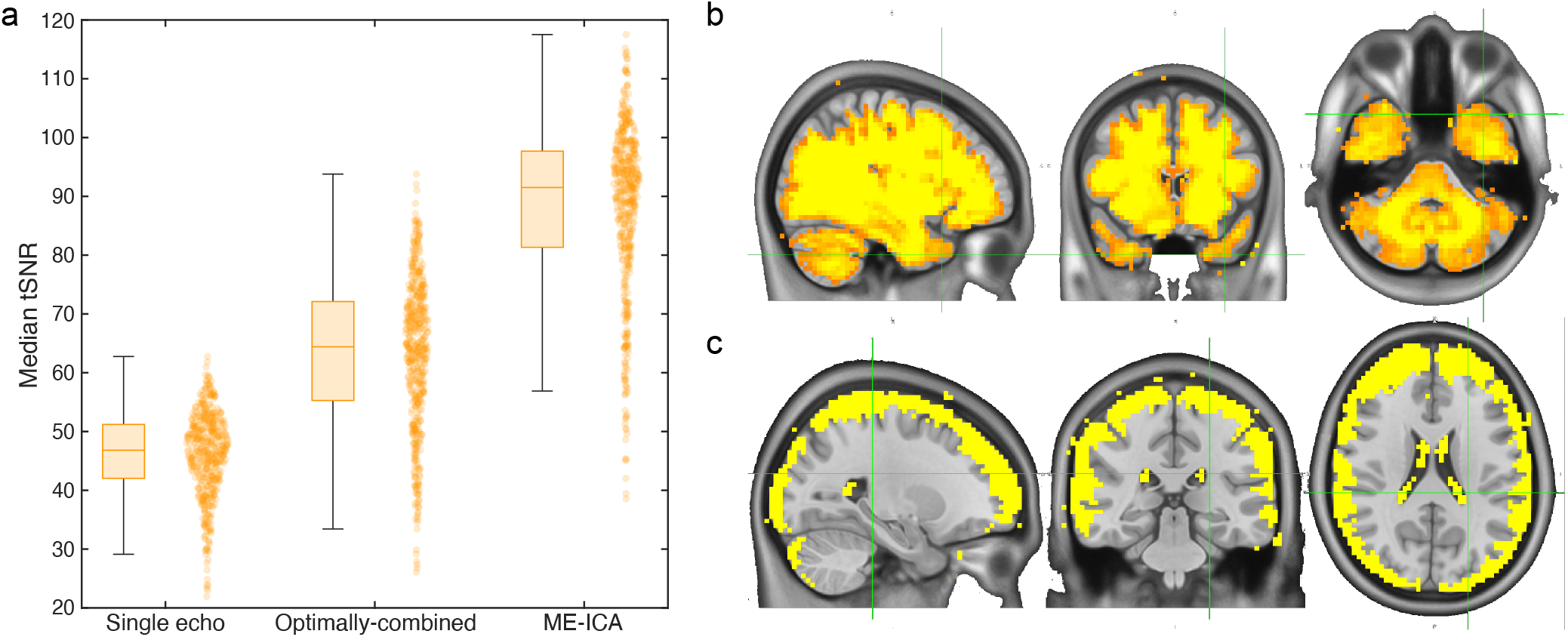
**a)** Comparison of temporal signal-to-noise ratio (tSNR) between single-echo data, optimally combined multi-echo data, and cleaned data via the multi-echo independent component analysis (ME-ICA). **b)** Spatial distribution of temporal signal-to-noise ratio (SNR) differences. Largest tSNR increase in the optimally combined multi-echo data compared to single-echo data. **c)** Further tSNR increase when using cleaned data via multi-echo ICA compared to the optimally combined data.

While we retained and released all the data, we used the above process to generate a table of recommended exclusions for users who do not wish to re-run a dedicated quality control analysis (see Supplementary Information Table 1). Among the functional MRI images, we excluded specific tasks for 7 participants, i.e., one or more among rest, meditation, music, or movie, as indicated in Fig. 2a. Among the anatomical MRI files, we excluded 8 structural T1w images from the psilocybin session but retained all the T1w images from the baseline session, as indicated in Fig. 2e. These are the exclusions that were applied in our exploratory analysis of the PsiConnect MRI data^45^.

### Cleaning the data removed slow drifts in the global signal

Functional MRI recordings often show a drift over time, due to magnetic field fluctuations caused by the scanner hardware, or by physiological factors such as slow variations in respiration or cardiac cycles that can introduce low-frequency fluctuations in the BOLD signal. This drift is evident in the raw data but is largely removed after cleaning via ME-ICA, and even more so after including white matter and SF signals as regressors in the GLM (see Supplementary Fig. 4). Since cleaning didn’t include global signal regression nor high-pass filtering, it is reasonable that some of these slow physiological drifts were captured by the confounding signals that were regressed.

**Figure 4.**
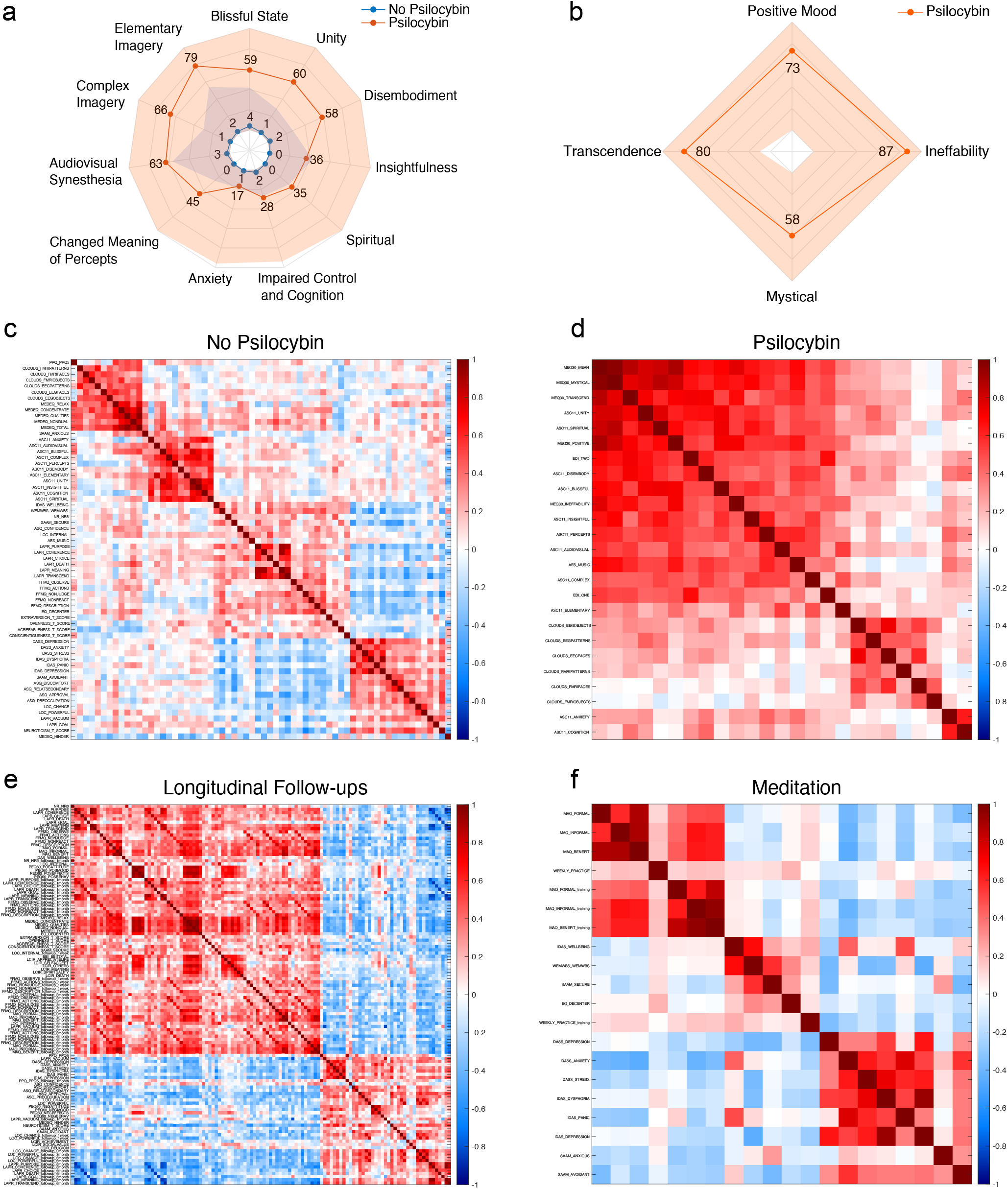
**a)** Radar plot of the 11-Dimension Altered States of Consciousness (ASC) Questionnaire, showing minimum-maximum range (shaded) and mean scores (solid line) across participants in both the psilocybin and no-psilocybin sessions. **b)** Radar plot of Mystical Experiences Questionnaire (MEQ30) showing minimum-maximum range (shaded) and mean scores (solid line) for 4 dimensions across participants. **c)** Pearson correlation between all pairs of behavioural scores acquired during baseline session (no psilocybin). **d)** Pearson correlation between all pairs of behavioural scores acquired after psilocybin administration. **e)** Pearson correlation between all pairs of behavioural scores acquired during longitudinal follow-ups. **f)** Pearson correlation between all pairs of behavioural scores related to the meditation scans and the 8-week meditation training (for half of the participants).

### Signal-to-noise ratio

The second echo was used as a proxy for single-echo data since the second echo time matches the typical value used in single-echo acquisitions (∼30 ms). The temporal SNR (tSNR) of this pseudo single-echo data was compared to the optimally combined multi-echo data and to the cleaned ME-ICA data, both at the subject- and at the voxel-level. For each scan, the median tSNR across all voxels is higher in the optimally combined multi-echo data than in the single-echo data, and it further increases after ME-ICA cleaning (Fig. 3a). This first result reveals the overall trend but ignores the spatial distribution of the tSNR across voxels. In order to locate these tSNR changes, we computed the median difference between tSNR values of each voxel (by subtracting the single-echo tSNR value from the optimally combined tSNR). The spatial maps shown in Fig. 3b demonstrate that the largest improvement is achieved in brain regions prone to signal dropout, including the temporal pole, ventromedial prefrontal cortex, and subcortical areas. In addition, Fig. 3c shows that the largest SNR improvement of ME-ICA over the optimally combined data is obtained in superficial grey matter voxels. Our results successfully replicate the median subject-level tSNR reported in^2^ and confirm that the benefits of the multi-echo data and ME-ICA procedures are region-specific. Importantly, the optimally combined multi-echo data shows higher tSNR in regions prone to signal dropout and in subcortical areas that are particularly relevant in the context of psychedelics. However, we must be careful not to understate the drawbacks of multi-echo fMRI sequences. The higher complexity in the acquisition and processing stages were mentioned above, as well as the reduced spatial resolution. In addition, the louder scanner noise could be distressing in psychedelic studies and might complicate task acquisitions involving complex auditory stimuli.

### Behavioral scores

The acute effects assessed using the 11-Dimension Altered States of Consciousness (11-D ASC) scale^8^ and the Mystical Experience Questionnaire (MEQ30)^46^ illustrate substantial group-level subjective effects intensity from the standardised 19mg dose of psilocybin (Fig. 4a,b). To provide an overview of the other behavioural scores available in the dataset, we present the matrices of pairwise Pearson correlation between the scores acquired during the psilocybin and no-psilocybin sessions, as well as during the meditation training and the longitudinal follow-ups (Fig. 4c,d,e,f). The matrices have been clustered to highlight sets of highly intercorrelated behavioural scores, which reflect key theoretical and conceptual distinctions in psychedelic effects. For example, the clusters identified during the psilocybin session (Fig. 4d) distinguish between: positively felt self- and boundary-dissolving effects, which characterise the high-level associative, abstract and existential experiential qualities (e.g., transcendence, bliss, unity); sensory-hallucinogenic effects (e.g., visual imagery, synaesthesia); and dysphoric effects (e.g., impaired control and cognition, and anxiety). We also note that negative effects, particularly anxiety, were infrequent or rated as low by the participants. Two clusters also emerge in the follow-up questionnaires and in the meditation-related scores (Fig. 4e,f), separating positive (purpose, meaning, coherence, well-being) and negative scores (stress, dysphoria, anxiety, discomfort).

### Usage Notes

Quality control of structural and functional data:

1. mriqc_singularity.sh (run MRIqc)
2. mriqc_PCA.m (obtain the plots in Fig. 2 based on the T1w and BOLD reports generated by MRIqc)

Pre-processing:

1. fmriprep_singularity.sh

Cleaning:

1. tedana.sh (identify non-BOLD independent components using ME-ICA)
2. glm_tedana_cleaning.m (fit a general linear model to regress head motion, cerebrospinal fluid, white matter, and non-BOLD independent components labelled as artifacts during the previous ME-ICA step. There is also the option to regress the global signal.)

## Code availability

Open-source code for all data analysis pipelines will be made available on GitHub upon publication (https://github.com/razilab). These pipelines used the following software packages: fMRIprep 22.0.2 (https://fmriprep.org), tedana 0.0.12 (https://tedana.readthedocs.io), MRIqc 22.0.6 (https://mriqc.readthedocs.io), MATLAB R2022a (https://www.mathworks.com), SPM12 (https://www.fil.ion.ucl.ac.uk/spm/software/spm12), Freesurfer 7.2 (https://surfer.nmr.mgh.harvard.edu), FSL v.6.0.7 (https://fsl.fmrib.ox.ac.uk/fsl/fslwiki), FieldTrip 20240916 (https://www.fieldtriptoolbox.org). LZ complexity was computed by adapting open-source code by Fernando Rosas and Pedro Mediano (https://information-dynamics.github.io/complexity/information/2019/06/26/lempel-ziv.html). Projection of volumetric fMRI data to the cortical surface was performed by adapting open-source code by Pang et al. available on GitHub (https://github.com/NSBLab/BrainEigenmodes).

## Acknowledgements

The authors acknowledge the facilities and scientific and technical assistance of the National Imaging Facility (NIF), a National Collaborative Research Infrastructure Strategy (NCRIS) capability at Monash Biomedical Imaging (MBI), a Technology Research Platform at Monash University. This research is funded by the Australian Research Council (Refs: DE170100128 and DP200100757) and the Australian National Health and Medical Research Council (Investigator Grant 1194910). A.R. is affiliated with The Wellcome Centre for Human Neuroimaging, supported by core funding from Wellcome [203147/Z/16/Z]. A.R. is a CIFAR Azrieli Global Scholar in the Brain, Mind & Consciousness Program. We also acknowledge USONA Institute for providing psilocybin through their “Investigatational Drug Supply Program”.

## Author contributions statement

**LN**: Methodology, Software, Validation, Formal analysis, Data Curation, Visualization, Writing - Original Draft. **DS**: Conceptualization, Methodology, Formal analysis, Investigation, Data Curation, Writing - Review & Editing, Project administration. **TB**: Data Curation, Writing - Review & Editing. **MDG**: Methodology, Software, Investigation, Data Curation, Writing - Review & Editing. **SD**: Conceptualization, Methodology, Writing - Review & Editing. **JJ**: Investigation. **JK**: Investigation, Writing - Review & Editing. **MLW**: Conceptualization, Investigation, Writing - Review & Editing. **AR**: Conceptualization, Methodology, Investigation, Writing - Review & Editing, Supervision, Project administration, Funding acquisition.

