## Supplementary Material for "PsiConnect: A Multimodal Neuroimaging Study of Psilocybin-Induced Changes in Brain and Behaviour"

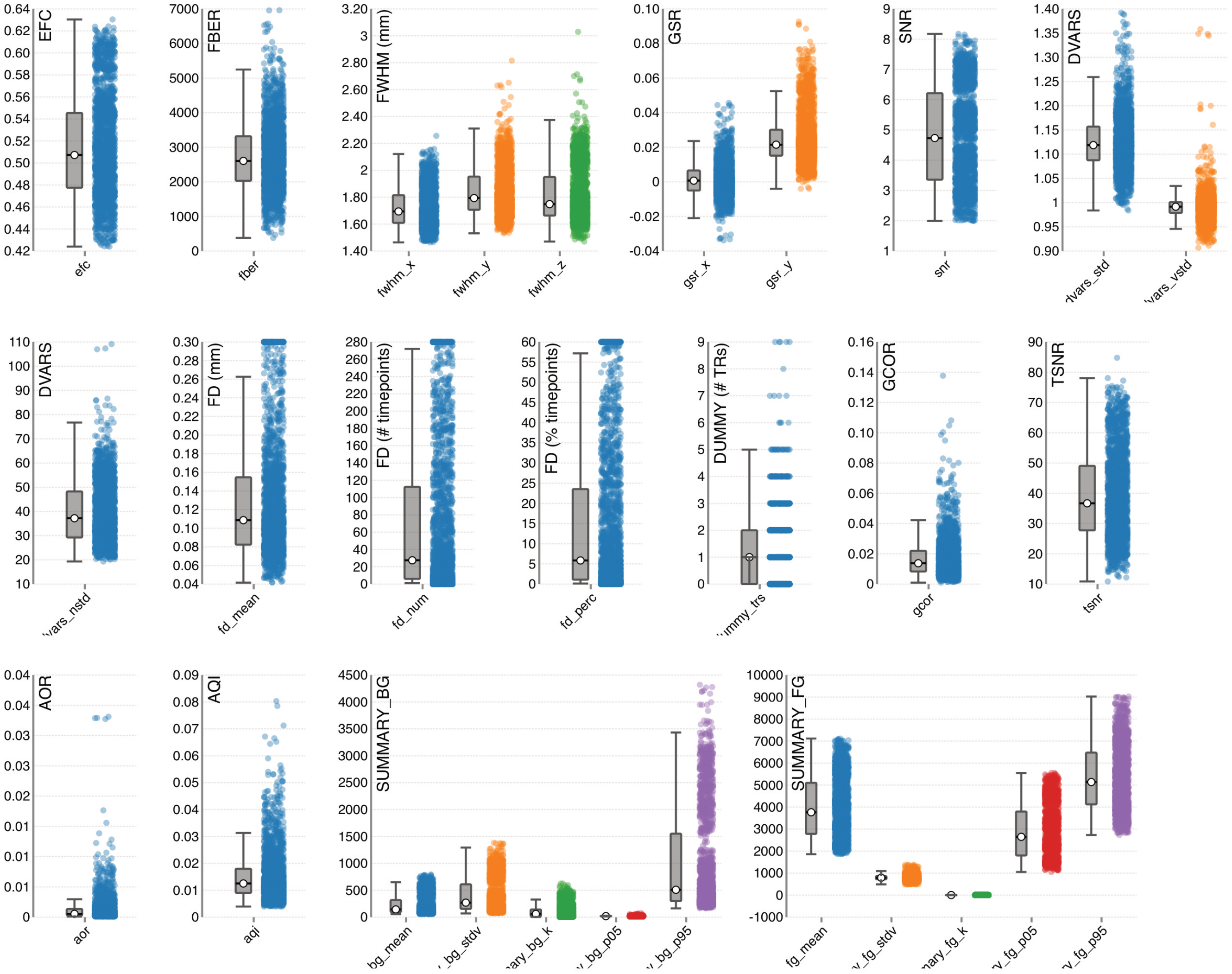


**Supplementary Fig. 1.** BOLD features computed via MRIQC.

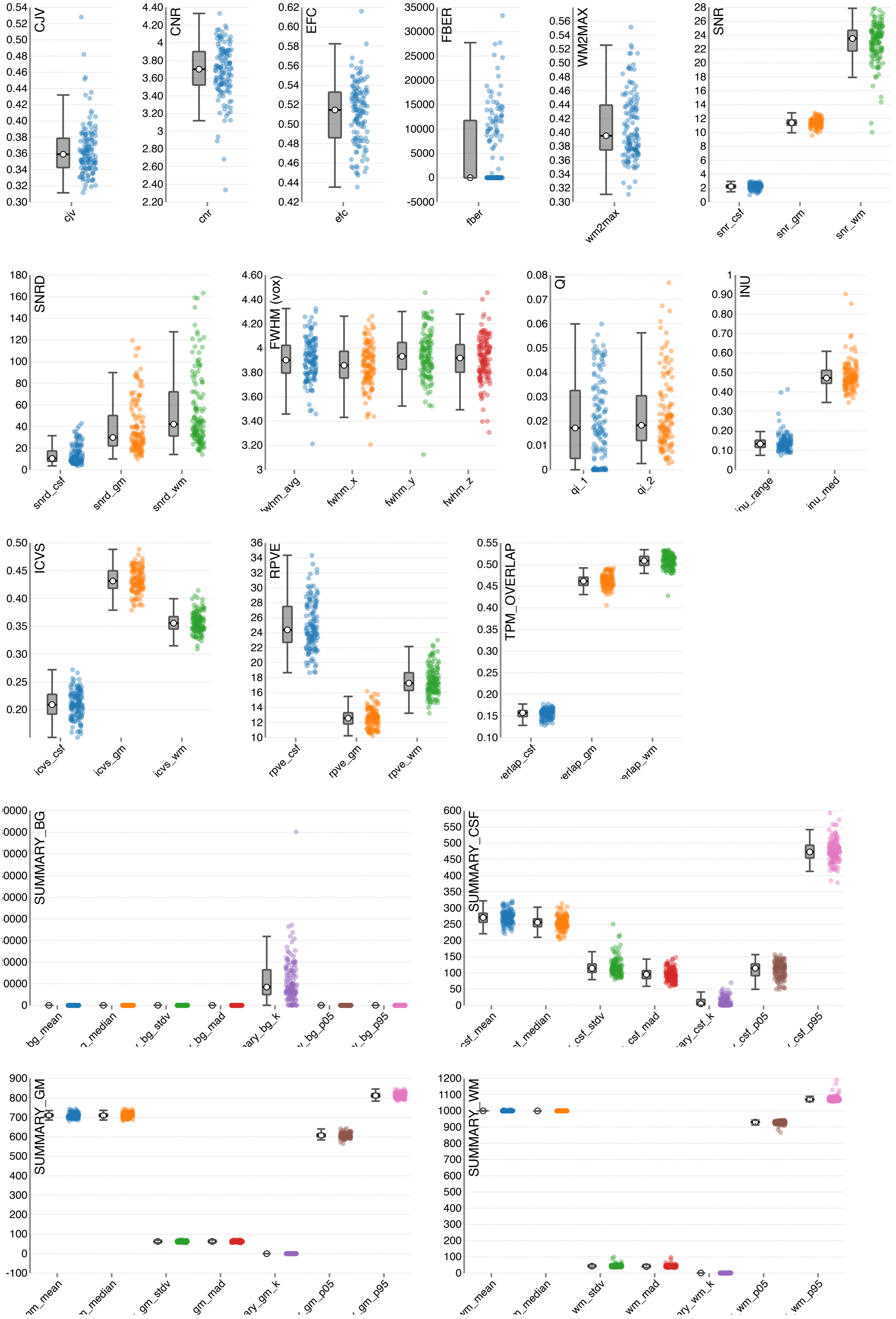


**Supplementary Fig. 2.** Anatomical (T1) features computed via MRIQC.

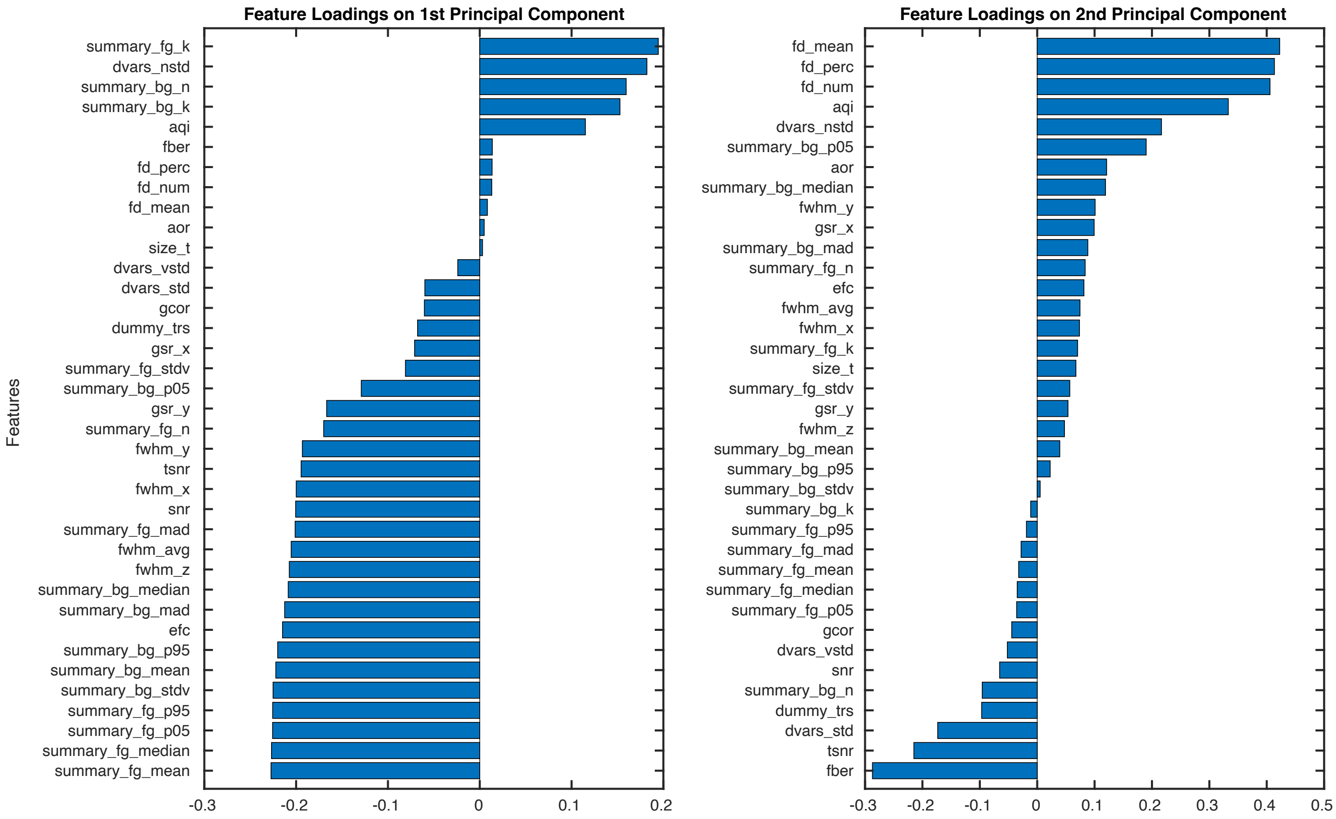


**Supplementary Fig. 3.** Principal component analysis (PCA) of the BOLD features computed via MRIQC. The two bar charts show the loading of each feature onto the first and second pricipal components. Head motion, quantified via the mean framewise displacement (FD) is the feature with the largest contribution to the second principal component. This explains why the outliers in Fig. 2a corresponded to participants who moved their head the most.

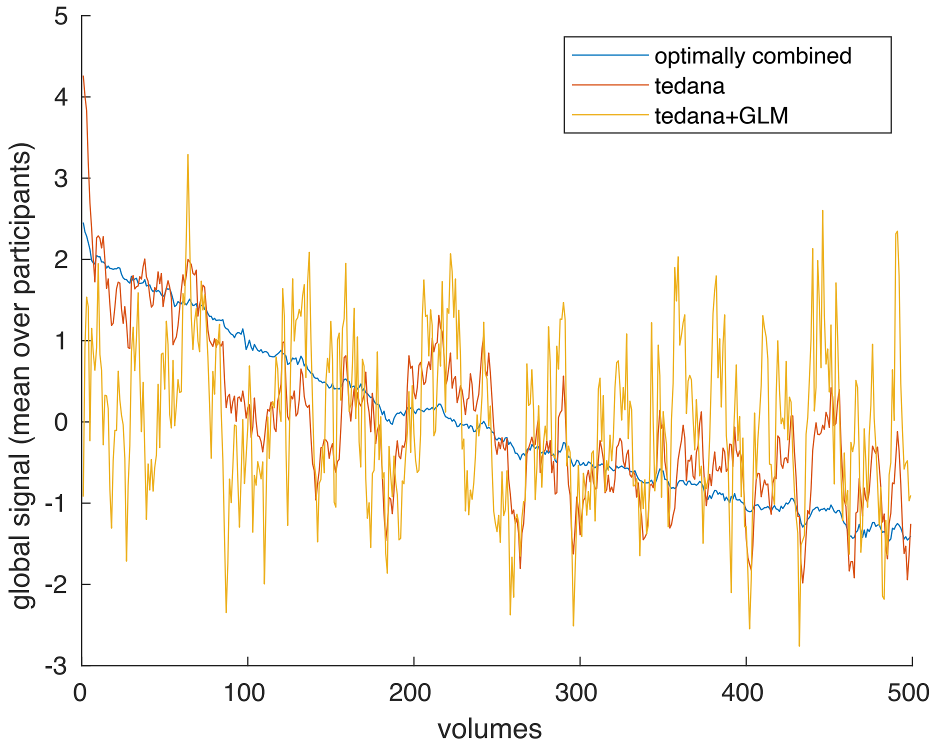


**Supplementary Fig. 4.** The slow drift in the global signal was removed by cleaning the data.

| Participant ID | Session | Modality | Task |
|---|---|---|---|
| PC003 | Psilocybin | BOLD | rest |
| PC003 | Psilocybin | BOLD | meditation |
| PC003 | Psilocybin | BOLD | music |
| PC003 | Psilocybin | BOLD | movie |
| PC005 | Psilocybin | BOLD | music |
| PC007 | Psilocybin | BOLD | rest |
| PC007 | Psilocybin | BOLD | meditation |
| PC007 | Psilocybin | BOLD | music |
| PC007 | Psilocybin | BOLD | movie |
| PC010 | Psilocybin | BOLD | music |
| PC022 | Psilocybin | BOLD | music |
| PC022 | Psilocybin | BOLD | movie |
| PC230 | Psilocybin | BOLD | movie |
| PC001 | Psilocybin | T1w | N/A |
| PC003 | Psilocybin | T1w | N/A |
| PC006 | Psilocybin | T1w | N/A |
| PC007 | Psilocybin | T1w | N/A |
| PC029 | Psilocybin | T1w | N/A |
| PC030 | Psilocybin | T1w | N/A |
| PC216 | Psilocybin | T1w | N/A |
| PC230 | Psilocybin | T1w | N/A |

**Supplementary Table. 1.** Suggested exclusions based on head motion and PCA analysis of BOLD MRIqc metrics.
